# Selective and Efficient Functionalization of P22 Virus-Like Particles Using an Asparaginyl Ligase

**DOI:** 10.64898/2026.07.02.736234

**Authors:** Maxim D. Harding, Mark A. Jackson, Kuok Yap, Pie Huda, David J. Craik, Frank Sainsbury, Nicole Lawrence

## Abstract

Protein cages provide useful scaffolds for nanoscale engineering due to their highly ordered structures and *in vivo* self-assembly. These scaffolds are amendable to late-stage conjugation, enabling expansion in functionality. However, many conjugation techniques either lack site-selectivity, require unnatural amino acid incorporation, or have bulky recognition motifs to facilitate ligation reactions. Here, an asparaginyl endopeptidase (AEP) enzyme with ligase activity is employed for the highly efficient functionalization of virus-like particles (VLPs) from *Salmonella* Typhimurium bacteriophage P22. The capacity of this enzyme to conjugate peptides and proteins onto assembled P22 VLPs under mild reaction conditions, via a minimal extension to the P22 coat protein C-terminus, is demonstrated. We extend the reaction efficiency to facilitate a one-pot dual-functionalization reaction whereby two therapeutically relevant receptor targeting domains are conjugated to P22 VLPs in a single step. Finally, we demonstrate the potential for AEP-mediated bioconjugation to bestow P22 VLPs with receptor-binding functionality *in vitro*. This work demonstrates the efficacy of AEP ligases as bioconjugation tools for site-selective functionalization of large molecular assemblies like VLPs.

## INTRODUCTION

Protein cages are nanoscale structures, comprised of one or more repeating subunits, that self-assemble to form a protein shell. Nature has independently evolved numerous protein cages, each with unique functionalities. Examples include viruses, ferritins, encapsulins, bacterial microcompartments, and the carboxysome^1^. The properties of these natural cages have enabled numerous biotechnological applications, including packaging of noncognate genes into adeno-associated viruses for gene delivery^2^, mineralization within ferritin^3^ or viral protein cages^4^, co-localizing heterologous enzymes within virus-like particles (VLPs) for metabolic engineering^5^, including constructing carbon fixing nanocages through rubisco encapsulation^6^. However, beyond capitalizing on the robust encapsulation capacity of natural protein cages, engineering functional domains to the exterior of these stable protein scaffolds can expand their possible applications^7^. Such modifications can be achieved at the DNA level through genetic fusion, or post-translationally via late-stage conjugation^7^. Genetic engineering of protein cages is limited in that the modified protein subunits must still enable organized self-assembly. Late-stage conjugation can overcome this hurdle by functionalizing already folded and assembled cage structures. Furthermore, late-stage conjugation can enable the production of complex fusion sequences and topologies inaccessible by genetic approaches.

Protein-mediated conjugation is a broadly accessible approach for achieving precise site-specific conjugation. For example, the SpyTag/SpyCatcher system enables highly specific isopeptide bond formation through a split *Streptococcus pyogenes* fibronectin-binding protein^8^. This approach has been successfully applied to functionalizing protein cages *in vitro* and *in vivo,* in both bacteria and plants^9–13^. However, a potential limitation of this system is that the bulky SpyTag/SpyCatcher sequence (12.7 kDa) remains incorporated in the final protein conjugate, which may be undesirable for some applications. Several enzymes with transpeptidase activity have also been used for late-stage conjugation to protein cages. The most common of these is sortase A, a transpeptidase from *Staphylococcus aureus* that facilitates amide bond formation between a five-residue C-terminal recognition sequence (LPXTG) and an incoming nucleophile with poly-glycine at the N-terminus^14^. By engineering this recognition sequence to the C-terminus of protein cage monomers, sortase A has been employed to conjugate desired functional moieties containing N-terminal poly-glycine tags^15, 16^, or vice versa^17^. Appropriate N- or C-terminal accessibility of the assembled monomers is the predominant factor that determines success of this approach; and therefore, protein cages with internalized termini are not amendable to sortase-mediated functionalization of assembled cages.

Recently, other conjugating enzymes have been employed for protein cage functionalization, including tyrosinase for bacteriophage MS2 VLPs^18^, transglutaminase for ferritin protein cages^19^, and small ubiquitin-like modifier enzyme Ubc9^20^, or ligase-type asparaginyl endopeptidase (AEP) for *Aa*LS protein cages^21^. Of these enzymatic approaches, AEP ligases combine a minimal recognition sequence with efficient reaction conditions that require minimal enzyme and low equivalents of the nucleophilic conjugation partner.

AEP ligases are a specialized subset of AEPs that evolved to preferentially cyclize peptide substrates by joining the N- and C-termini of peptides that contain a C-terminal tripeptide recognition sequence (commonly NGL)^22^. This recognition sequence is bound and cleaved by the AEP between the asparagine and glycine, forming an acyl-enzyme intermediate, which is then resolved by nucleophilic attack of an incoming N-terminal amine^23^. Multiple studies that explored the capabilities of AEP ligases in protein engineering applications have demonstrated the promiscuity of incoming amine nucleophiles and the successful conjugation of a range of peptide and protein substrates^21, 24–28^. Compared to other enzymes with ligase functionality, AEP ligases can be used at lower enzyme-to-substrate ratios, at ambient temperatures, and have a minimal carryover of recognition sequence following conjugation. Despite the interest in AEP ligases as protein engineering tools, their utility as highly efficient tools for functionalization of large molecular assemblies, such as protein cages, has been underexplored.

*Salmonella* Typhimurium bacteriophage P22 VLPs are versatile scaffolds for protein engineering. Comprised of 420 coat proteins (CP) and 100-330 scaffold proteins (SP) per VLP, CP and SP co-expression in *Escherichia coli* or *Nicotiana benthamiana* enables formation of 58 nm icosahedral protein cages^13, 29^. P22 VLPs are highly programmable, capable of both protein encapsulation through SP genetic fusion^29–31^, and external protein display achieved by fusion to externalized CP C-termini^15, 32^. Late-stage external protein display on P22 VLPs through the CP C-terminus has previously been explored by incorporating SpyTag for *in vitro* functionalization with SpyCatcher-tagged fusion proteins^32^, or the sortase A recognition sequence to facilitate enzyme-mediated functionalization^15^.

The ability to functionalize protein cages via a minimal proteinogenic recognition tag is highly desirable for modular late-stage engineering of protein cages. Here, we explore the use of [C247A]*Oa*AEP1, a catalytically optimized variant of the AEP ligase from *Oldenlandia affinis*^33^ hereafter referred to simply as AEP, for mediating late-stage functionalization of P22 VLPs with peptide and protein conjugation partners. To facilitate conjugation, the AEP recognition motif was added to the C-terminus of the P22 coat protein (CP), which had no impact on VLP assembly. Site-selective conjugation was explored using a cell-penetrating peptide, a fluorescent reporter protein, and two therapeutic receptor targeting domains. Finally, we demonstrate the capacity for P22 VLPs functionalized with these targeting domains to bind their cognate receptors *in vitro*.

## RESULTS AND DISCUSSION

### Modification of P22 VLP C-Terminus

To assess the feasibility of AEP-mediated functionalization of P22 VLPs, we first investigated the effect of inserting the AEP recognition sequence onto the P22 CP C-terminus on VLP assembly. We have previously demonstrated that AEP transpeptidation efficiency is improved by nickel ion quenching of the C-terminal recognition sequence when a histidine is included at the C-terminus (Figure 1A)^26^. Therefore, we included the optimized NGLH recognition sequence at the CP C-terminus, separated by a short flexible linker (GGSGG, see Table S1). This engineered CP, termed CP-NGLH, was co-expressed with the red fluorescent protein mRUBY3 fused to the C-terminus of P22 scaffold protein (SP) (SP-mRUBY3, Figure 1A, Table S1). Co-expression of both CP-NGLH and SP-mRUBY3 yielded correctly assembled VLPs with SP-mRUBY3 packaged, as determined by size-exclusion chromatography (SEC), sodium dodecyl-sulfate polyacrylamide gel electrophoresis (SDS-PAGE), and negative stain transmission electron microscopy (TEM) (Figure 1B, C, D). Further assessment of purified CP-NGLH by in-gel tryptic digest liquid chromatography tandem mass spectrometry (LC/MS-MS) confirmed the presence of the inserted C-terminal AEP recognition sequence (Table S2). These data confirm that addition of the AEP recognition sequence to the C-terminus of the P22 CP does not inhibit VLP assembly, enabling AEP-mediated transpeptidation to be investigated.

**Figure 1.**
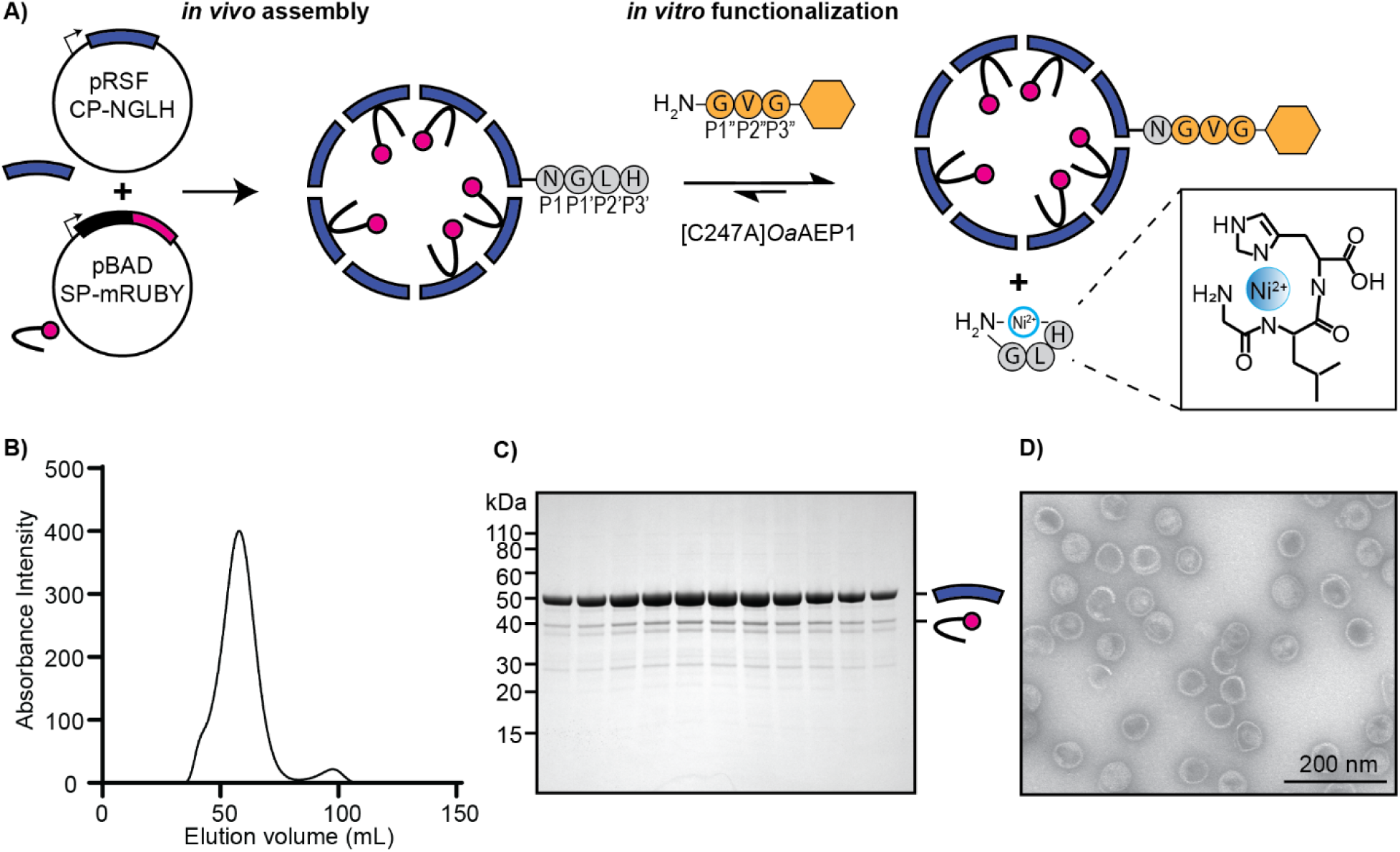
Engineering an AEP-recognition site onto the C-terminus of P22 coat protein (CP). A) Cartoon depicting the plasmids used for P22 VLP expression that lead to *in vivo* assembly in *E. coli,* and the transpeptidation reaction process (the GGSGG linker preceding the NGLH recognition sequence is not shown). Purified P22 VLPs with mRUBY3 cargo and engineered CP C-termini for AEP recognition are functionalized *in vitro* by the [C247A]*Oa*AEP1 enzyme which catalyzes the transpeptidation reaction. This enzyme cleaves between the asparagine and glycine amino acids (P1 and P1’, respectively). The optimized C-terminal leaving group sequence (GLH) is effectively quenched by Ni_2_+ in solution^26^. B) Chromatogram from size exclusion chromatography (SEC) of ultracentrifuged protein extracts from co-expression with CP-NGLH and SP-mRUBY3. The large peak beginning at 50 mL elution volume corresponds to P22 VLPs. C) Coomassie stained SDS-PAGE of fractions from SEC. The CP-NGLH species is the strong band at 50 kDa and SP-mRUBY3 is the band below at 40 kDa. D) Negative stain TEM of SEC purified and pooled P22 VLPs with C-terminal NGLH.

### Conjugation of a Synthetic Peptide

For proof-of-concept assessment for AEP-mediated conjugation to P22 VLPs, a membrane-selective cell-penetrating peptide was chosen. PDIP, a helix-turn-helix macrocyclic peptide, can target and enter malaria-infected red blood cells^34^ and cancer cells^35^, and its cell-penetrating properties have been successfully exploited through application as a peptide-conjugate for targeted delivery of antimalarial drugs^36, 37^, and anticancer peptides^38^ or drugs^39, 40^ *in vitro*. As a positively charged membrane-active peptide, PDIP is challenging to produce biosynthetically due to its inherent toxicity when accumulated in host expression cells^30^. Indeed, genetic fusion of PDIP to the CP C-terminus for external display failed to produce purifiable P22 VLPs when expressed in *E. coli*. Therefore, late-stage functionalization with chemically synthesized PDIP was evaluated, using a modified PDIP analogue with an N-terminal GVG sequence (herein referred to as GVG-PDIP) (Figure S1), as including a short hydrophobic residue (e.g. valine) at the P2’’ position (Figure 1A) is expected to increase AEP-transpeptidation efficiency^41^.

The benchmark for enzyme-mediated conjugation to the externalized C-terminus of the CP on P22 VLPs is sortase A^15^. Reactions using sortase A were previously shown to be optimal when two equivalents of enzyme to substrate were used, at a temperature of 42 °C for at least 6 h^15^. Here, to demonstrate the hypothesized advantage of using an AEP ligase, significantly milder reaction conditions were evaluated for the conjugation of GVG-PDIP to engineered P22 VLPs (Figure 2A). Reactions were performed with 15-fold less enzyme than VLP (CP-NGLH) substrate, with 4-equivalents of GVG-PDIP to VLP (CP-NGLH), at room temperature (22 °C) for a maximum of 2 h. Successful conjugation was assessed by quenching the reactions by dilution in 1% (v/v) formic acid at set time points and visualizing product formation by SDS-PAGE (Figure 2B). After 15 min of reaction, a higher molecular weight protein band could be seen by SDS-PAGE, which increased slightly over the 2 h reaction time (Figure 2B). Visualization of the reaction after 2 h by negative stain TEM confirmed the maintained structure of P22 VLPs (Figure 2B). The predicted importance of the N-terminal GVG extension to PDIP was confirmed by repeating the conjugation reaction with PDIP without this extension (Figure 2C). Even after 16 h incubation, a CP-PDIP conjugation product was almost undetectable by SDS-PAGE (Figure 2C) but was present as confirmed by western blot analysis (Figure S2). This poor transpeptidation efficiency contrasts with the reaction with GVG-PDIP for 16 h, which yielded VLPs with 35% of the total CP functionalized with GVG-PDIP (Figure 2D, Figure S2). Given the macrocyclic structure of PDIP, constrained by a disulfide bond between N- and C-terminal cysteine residues (Figure 2C), it is not surprising that an N-terminal extension is required to achieve efficient conjugation of PDIP to CP-NGLH on P22 VLPs.

**Figure 2.**
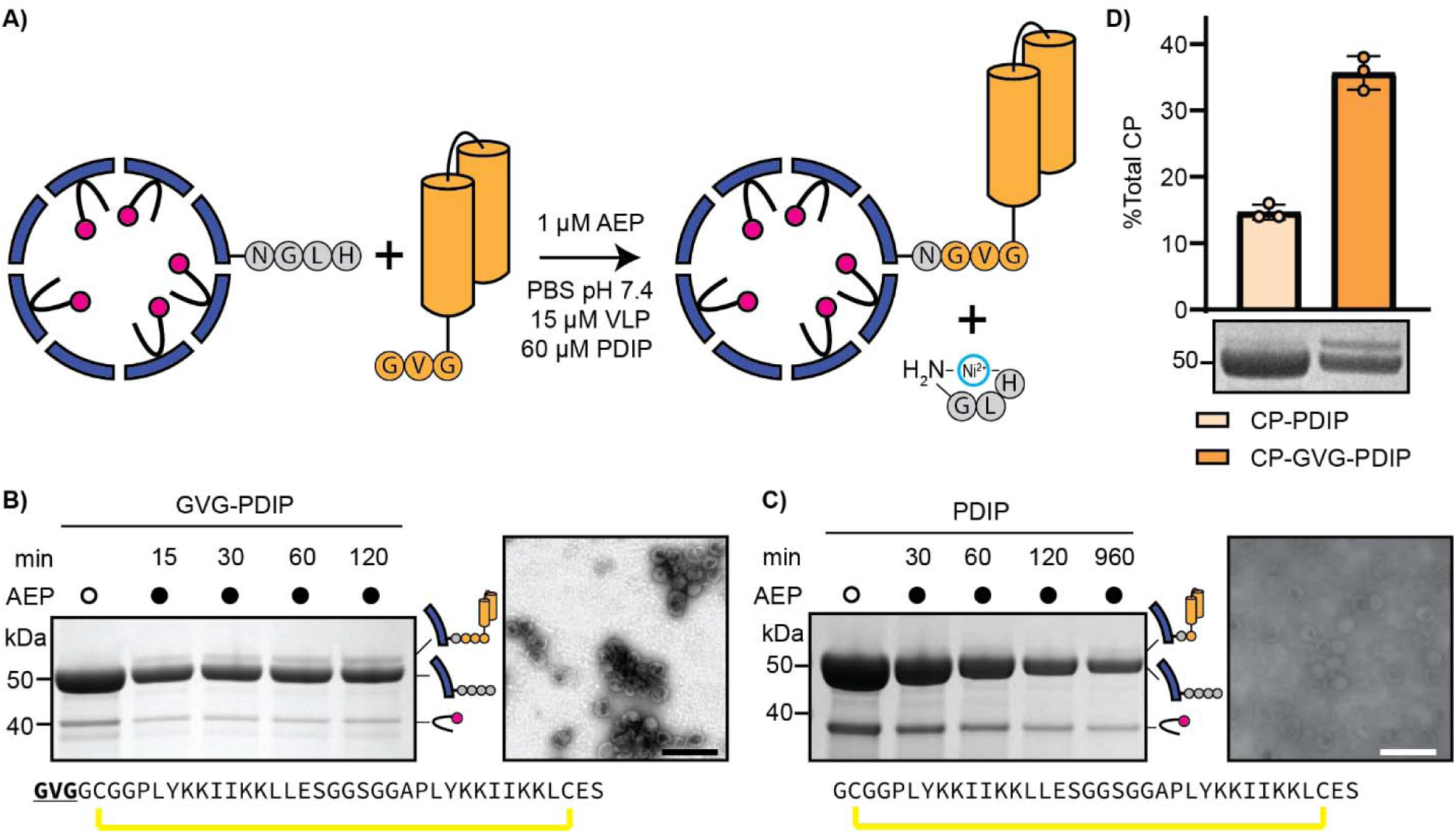
AEP-mediated conjugation of a synthetic peptide. A) Illustration of AEP-mediated conjugation of GVG-PDIP to assembled P22 VLPs (with mRUBY3 cargo) via CP-NGLH. Reaction conditions are shown. B) Coomassie stained SDS-PAGE of quenched reaction time points (shown in minutes) and a no-enzyme control. Negative stain TEM of conjugated P22 VLPs following 2 h incubation (scale bar indicates 200 nm). Amino acid sequence of GVG-PDIP with N-terminal GVG extension underlined and bolded. Yellow lines indicate disulfide bonds. Protein species are illustrated on the right side of the gel. C) Data shown as in B) but for PDIP without N-terminal GVG extension. D) Coomassie stained SDS-PAGE gel showing CP-NGLH following 16 h conjugation with either PDIP or GVG-PDIP (left and right respectively), with bar graph illustrating densitometry-based quantification of PDIP or GVG-PDIP conjugated CP (top band) as a percentage of total CP (top + bottom bands).

Evaluation of the GVG-PDIP conjugation reaction following 16 h incubation by negative stain TEM revealed large clumps of VLPs (that were not observed after 2 h), with individual VLPs being almost indistinguishable by TEM (Figure S3A). A scaled 16 h reaction (1 mL volume total, with same equivalents as aforementioned) displayed distinct turbidity compared to a no-enzyme control (Figure S3B). Centrifugation of the diluted reaction produced a pink pellet (Figure S3B, C) that suggested aggregation of the mRUBY3-containing VLPs. Because GVG-PDIP contains multiple lysine residues, we considered whether the recently reported capability of AEP-mediated lysine isopeptide bond formation^42^ could be contributing to VLP crosslinking. However, SDS-PAGE analysis of the aggregating material did not reveal the presence of higher molecular weight oligomers that would be expected for CP-GVG-PDIP-CP covalent linkages (Figure S3D). The absence of oligomers is not consistent with observations of P22 VLP aggregation caused by sortase-mediated GFP crosslinking^15^, suggesting that changes in the overall surface characteristics of GVG-PDIP-functionalized VLPs drove aggregation via non-covalent interactions. Importantly, this aggregation was not observed with other domains of interest (see below), suggesting that this phenomenon is not an inherent limitation of AEP-mediated P22 VLP functionalization. Having demonstrated proof-of-concept for AEP-mediated functionalization to P22 VLPs with a small helical peptide, we next evaluated this approach with recombinant globular proteins.

### Application to Protein Conjugation

To further evaluate the versatility of AEP-mediated late-stage functionalization, we sought to conjugate larger nucleophiles onto P22 VLPs. For this purpose, we examined the efficiency of AEP-mediated transpeptidation of a 29 kDa protein – a super folder green fluorescent protein containing an N-terminal GL extension (GL-sfGFP) that was produced in *E. coli* (Figure S4). The reported conjugation of GFP to P22 VLPs using sortase A employed 10 equivalents of GFP and 2 equivalents of sortase A relative to the P22 CP^15^. By contrast, the AEP-mediated conjugation reaction tested here employed 15-fold less enzyme relative to CP-NGLH, with only 4-fold excess GL-sfGFP (Figure 3A). Assessment of reaction progression by SDS-PAGE at 15, 30, 60, and 120 min time points demonstrated efficient GL-sfGFP conjugation onto P22 VLPs over time (Figure 3B), with the ∼80 kDa protein band identified by tryptic digest LC-MS/MS to be correctly conjugated CP-NGL-sfGFP (Table S2). Assessment of the 2 h reaction by negative stain TEM also confirmed the maintenance of P22 VLP structure following GL-sfGFP conjugation (Figure 3D). Quantification of this reaction over time by densitometry analysis revealed an average of 32% of total P22 CP being conjugated with GL-sfGFP after 2 h (Figure 3C, Figure S5), corresponding to 134 molecules of GL-sfGFP per P22 VLP (assuming 420 CP per VLP). The amount of conjugated GL-sfGFP is similar to reports using sortase A, with approximately 140 GFP molecules per VLP achieved^15^. However, the AEP-mediated approach presented here achieved this level of conjugation with 30-equivalents less enzyme, 6-equivalents less GL-sfGFP nucleophile and in only 2 h at room temperature.

**Figure 3.**
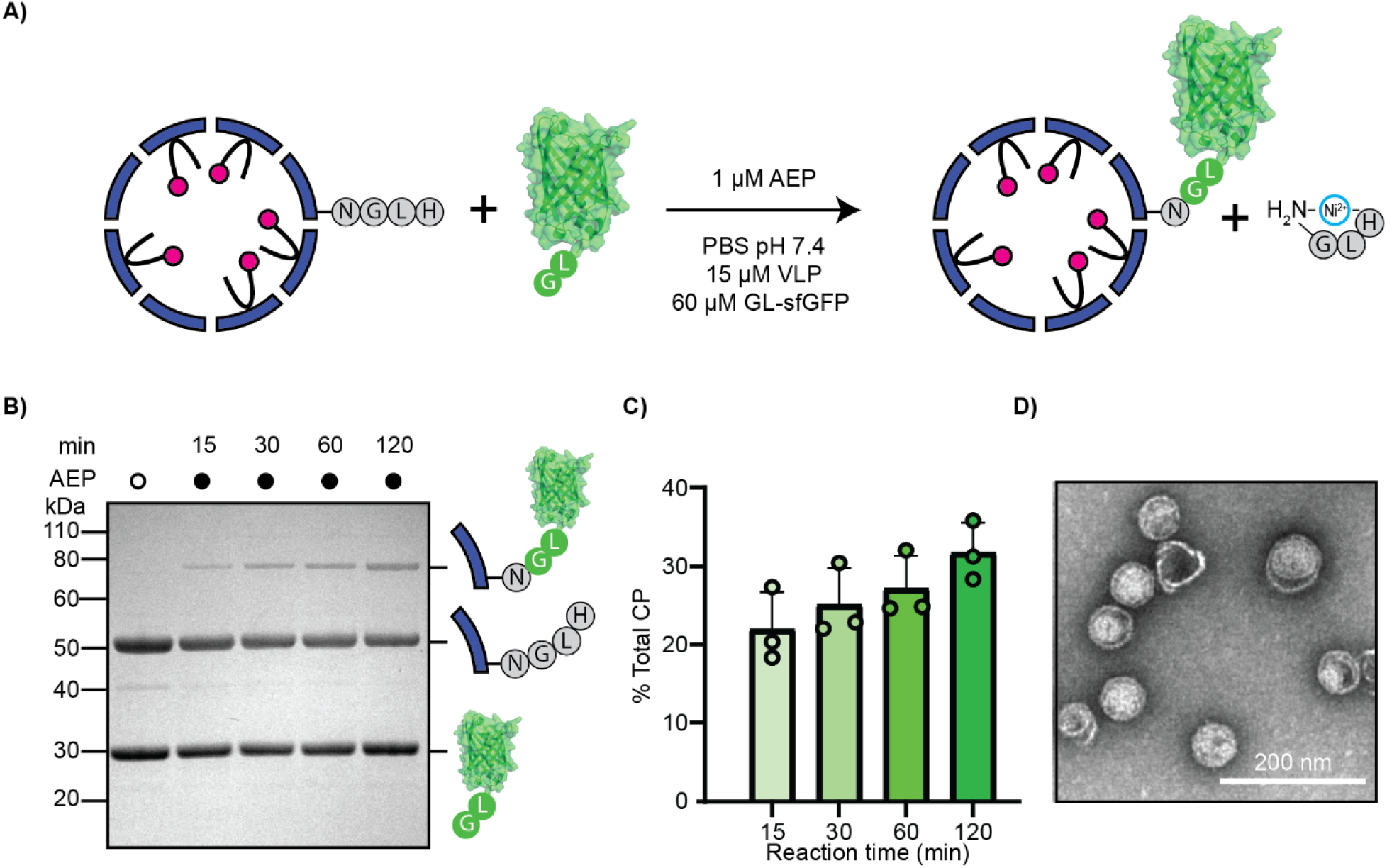
Conjugation of superfolder (sf) GFP to P22 VLPs. A) Illustration of the AEP-mediated conjugation reaction. B) Representative Coomassie stained SDS-PAGE of quenched reaction time points (time indicated in min) and a no enzyme control. Representations of each major protein species present are displayed on the right side of the gel. C) Densitometry-based quantification of the GL-sfGFP conjugation efficiency, represented as a percentage of the total CP (n=3 technical repeats). D) Negative stain TEM of GL-sfGFP conjugated P22 VLPs after 2 h incubation. Scale bar indicates 200 nm.

AEP-mediated conjugation of both peptide (GVG-PDIP) and protein (GL-sfGFP) nucleophiles resulted in a maximal proportion of ∼ 30% conjugated CP. This apparent limit in percentage of total VLP CP able to be functionalized is probably due to steric restriction caused by proximity of the CP C-termini at both pentameric and hexameric VLP assembly interfaces (Figure S6), but altered attraction of incoming nucleophiles due to changes in overall VLP surface characteristics may also contribute. This outcome is consistent with previous studies that explored functionalization of P22 VLPs via sortase A and SpyTag/SpyCatcher^15, 32^. P22 CPs assemble into 60 hexamers (see hexamer structure in Figure S6) and 12 pentamers to form a P22 VLP. Within this structural arrangement, the C-termini of adjacent CPs are approximately 30Å apart. Given GFP has dimensions of approximately 30×40Å, it is reasonable to assume that conjugation to one CP may preclude conjugation to adjacent CPs.

### Functionalization of P22 VLPs with Receptor-Targeting Domains

Having demonstrated the ability to conjugate both peptides and larger proteins to P22 VLPs late-stage, we next explored the capacity for functionalization with therapeutically relevant targeting domains. Engineering VLPs and other protein cages as payload delivery vehicles for gene editing, mRNA vaccine delivery, and cancer therapy is an exciting area of therapeutics development^43–45^. We propose that AEP-mediated functionalization of VLPs with target specific molecules can drive receptor binding and selective delivery of encapsulated payloads to tissues and cells of interest, an important requirement for therapeutic development of engineered VLPs and other protein cages. As a proof of concept, the human epidermal growth factor receptor 2 (HER2) and epidermal growth factor receptor (EGFR) were explored as clinically relevant cancer receptors^46^, which we targeted by functionalizing P22 VLPs with ZHER2 affibody and 9G8 nanobody respectively^47, 48^ (Figure 4A). Both these targeting domains were expressed in *E. coli* and processed to leave an N-terminal GVG extension, creating GVG-ZHER2 and GVG-9G8 (Figure S7 and S8). AEP-mediated conjugation reactions were performed as described for GL-sfGFP above to successfully produce VLPs that were functionalized with GVG-ZHER2 or GVG-9G8 respectively, as demonstrated by SDS-PAGE and tryptic digest LC-MS/MS (Figure 4B, 4C, Table S2). The conjugation efficiency between GVG-ZHER2 and GVG-9G8 differed, with GVG-ZHER2 achieving over 30% total CP functionalization after 3 h compared to GVG-9G8 which achieved 20% as determined by densitometric analysis (Figure 4E, 4F). TEM imaging of P22 VLPs following 2 h reaction confirmed that correct morphology was maintained for VLPs functionalized with GVG-ZHER2 or GVG-9G8 (Figure 4H, 4I).

**Figure 4.**
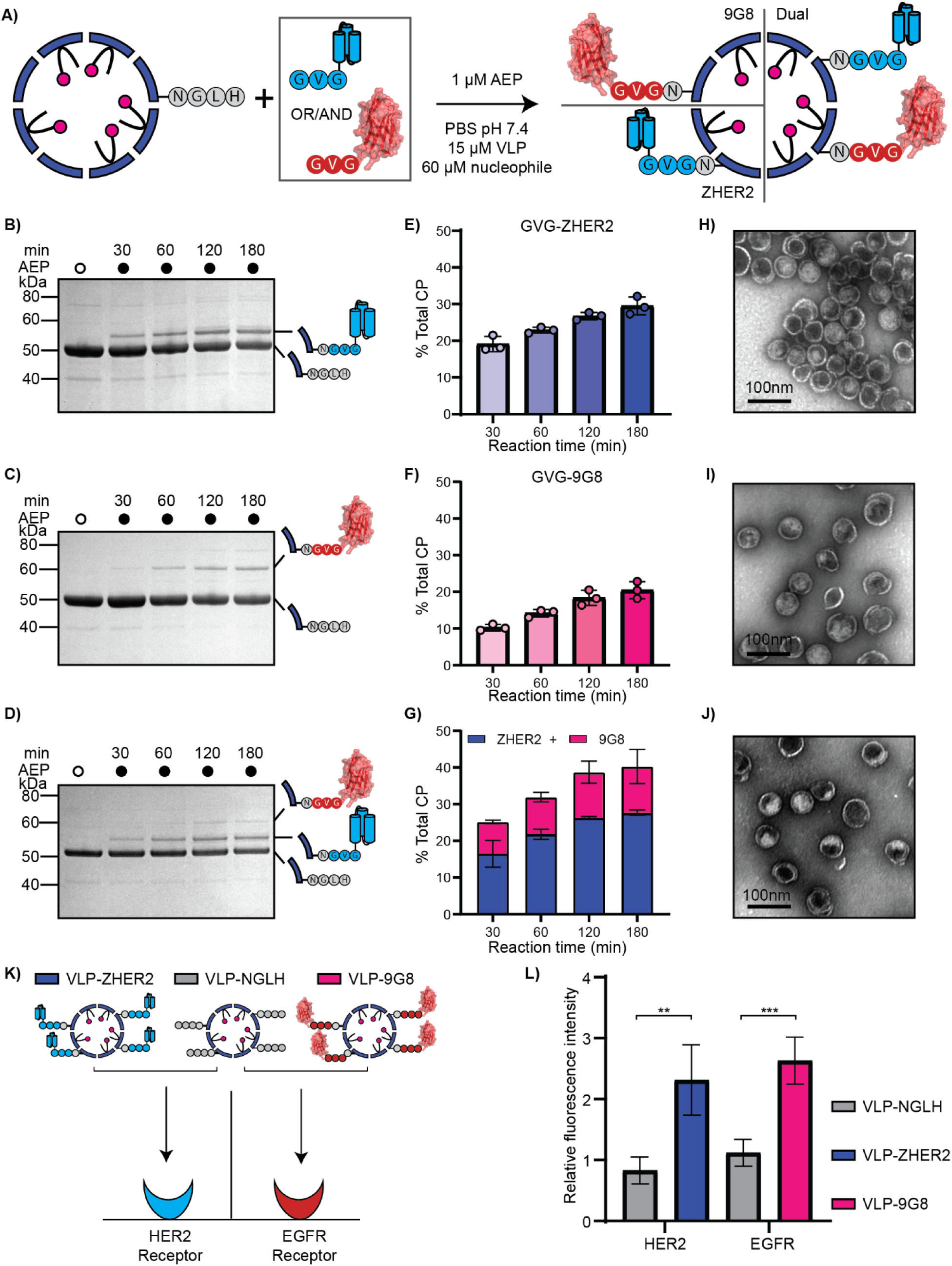
Functionalization of P22 VLPs with receptor-targeting domains. A) Illustration of the bioconjugation reaction, indicating reaction conditions. Here, both the conjugation of each receptor-targeting domain alone and a one-pot reaction with both domains was evaluated. Coomassie stained SDS-PAGE of reaction with GVG-ZHER2 (B), GVG-9G8 (C), and a one-pot reaction with both GVG-ZHER2 and GVG-9G8 (D) over time (min). Densitometric quantification for reactions over time for GVG-ZHER2 (E), GVG-9G8 (F), and one-pot reaction with both (G) (n=3 technical repeats each). Negative stain TEM images for VLPs functionalized with (H) GVG-ZHER2, (I) GVG-9G8, (J) and both. K) Illustration of functionalized VLPs and VLP control assessed for *in vitro* binding to HER2 and EGFR receptors. L) Relative fluorescence intensity (corresponding to VLP-encapsulated mRUBY3 fluorescence) for VLP incubation with HER2 or EGFR receptors. Data is shown as relative mean fluorescence intensity ± SD from a single experiment with 5 technical replicates for each VLP/receptor pair. The data was normalized to account for any nonspecific binding to BSA and/or receptor. Functionalized VLP vs VLP control were analyzed for each receptor with a Welch’s t test (** *p* < 0.01, *** *p* < 0.001) using GraphPad Prims v10.

To determine whether AEP-mediated conjugation could be used to produce dual-functionalized VLPs, like the demonstrated co-functionalization to protein cages using SpyTag/SpyCatcher^49^, we performed a one-pot conjugation reaction with both GVG-ZHER2 and GVG-9G8. We used the same concentration of AEP and VLP as for the single nucleophile reactions above, but used only 30 µM each of GVG-ZHER2 and GVG-9G8, to keep the total nucleophile ratio at 4 molar equivalents of the VLP (Figure 4A). Reaction monitoring by SDS-PAGE confirmed the ability for AEPs to mediate one-pot dual functionalization, with CP-GVG-ZHER2 and CP-GVG-9G8 both visualized by SDS-PAGE (Figure 4D). Unsurprisingly given the disparity in conjugation efficiency between GVG-ZHER2 and GVG-9G8 alone, more GVG-ZHER2 was conjugated than GVG-9G8, accounting for approximately 70% of the functionalized CP (Figure 4G). Densitometric analysis of CP functionalization for the one-pot reaction was calculated to be ∼40% of total CP after 2 h incubation, which is the highest of the reactions tested here but within range of the expected maximal level for P22 CP functionalization^15, 32^. One-pot dual-functionalized P22 VLPs, like all VLPs functionalized by AEP-mediated transpeptidation in this study, maintained correct morphology as shown by TEM (Figure 4J).

To confirm that the ZHER2 and 9G8 receptor targeting domains bind to their respective receptors when conjugated to P22 VLPs, we performed an *in vitro* receptor-binding assay and examined affinity of VLP-ZHER2 and VLP-9G8 for recombinantly produced HER2 and EGFR receptors compared to an unfunctionalized control VLP (CP-NGLH) (Figure 4K). For both VLP-ZHER2 and VLP-9G8, the relative fluorescence intensity when incubated with their respective cognate receptors were significantly greater than for the VLP (CP-NGLH) control (Figure 4L). This data confirms that GVG-ZHER2 and GVG-9G8 retain their receptor binding capability when conjugated to P22 VLPs by AEP enzymes *in vitro,* providing proof-of-concept for applying AEP-mediated conjugation to functionalize protein cages like P22 VLPs with therapeutically relevant proteins domains.

## CONCLUSIONS

Late-stage modification is a powerful tool for enhancing the functionality of proteins of interest, including complex multimeric assemblies like VLPs. AEP ligases are highly effective protein engineering tools, capable of rapid, site-selective conjugation of peptides and proteins, with minimal quantities of enzyme required. Here, we explored the use of an AEP ligase for late-stage functionalization of P22 VLPs *in vitro*. By including the minimal AEP recognition sequence (NGLH) to the C-terminus of the P22 CP, AEP ligases can effectively functionalize P22 VLPs with peptides or proteins containing a range of tolerated N-terminal sequences, that can be modified to enhance conjugation efficiency (e.g. varying linker length, presence of short hydrophobic residue at P2”)^41^. We demonstrated the efficacy of AEP-mediated functionalization for peptides and proteins ranging in size from 4 to 28 kDa, which was expanded to include one-pot dual-functionalization of P22 VLPs with two receptor targeting domains, ZHER2 affibody (7 kDa) and 9G8 nanobody (14 kDa). Notably, VLPs functionalized with these targeting domains bound to their respective HER2 and EGFR receptors in an *in vitro* receptor-binding assay.

The approach demonstrated here is suited to functionalizing protein cages with externalized CP C-termini; however, future work could expand the scope of protein cage engineering by employing AEP ligases for conjugation through exposed lysine residues within the CP sequence, a capability that we have recently demonstrated for other proteins^42, 50^. Additionally, to allow a greater degree of control over dual functionalization reactions, future work could employ two recently engineered AEP variants with orthogonal recognition sequences^28^ to facilitate sequential transpeptidation of different nucleophiles. An additional study that explored applications of AEPs for protein-protein bioconjugation reactions in *E. coli*^51^ suggests the potential for also using AEPs to facilitate *in vivo* functionalization of P22 VLPs, which given the ability to assemble P22 VLPs in plants^13^, may be expanded to *in planta* protein engineering applications in the future^52^. Ultimately, the work presented here demonstrates that the efficiency and durability of AEP ligases, which has made them favored tools for protein engineering, can be applied for functionalizing large protein assemblies such as P22 VLPs, thereby opening future possibilities for creating newly functionalized VLPs which are inaccessible by genetic fusion approaches.

## METHODS AND MATERIALS

### Molecular Cloning

Plasmids for VLP expression (pRSF CP and pBAD SP-mRUBY3) were cloned as described previously^31^. C-terminal modification to the P22 CP was achieved by inverse PCR on the pRSF CP plasmid to insert the additional C-terminal sequence followed by plasmid re-ligation using New England Biolabs (NEB) HiFi assembly Master Mix. GVG-ZHER2 and GVG-9G8 were ordered as dsDNA gene blocks from IDT DNA Technologies and cloned into a modified pET20b(+) plasmid (containing no pelB periplasmic signal peptide) by Gibson assembly using HiFi assembly Master Mix (New England Biolabs). GL-sfGFP was gifted as a glycerol stock of BL21(DE3) cells by Dr Yan Zhou (University of Queensland). All cloned plasmids were sequenced by whole plasmid sequencing (Plasmidsaurus) before transforming into chemically competent *E. coli* BL21(DE3) cells.

### P22 VLP Expression and Purification

P22 VLP assembly was achieved through co-expression of pRSF CP-NGLH and pBAD SP-mRUBY3, harboring kanamycin and ampicillin selection cassettes respectively, with induction at A600 of 0.6 with both 1 mM IPTG and 0.02 % (v/v final concentration) L-Arabinose. Following induction, cultures were incubated at 28 °C with shaking at 220 rpm. Following 16 h expression, cells were harvested by centrifugation at 4,500 g, resuspended in PBS (pH 7.4) and lysed by three passages through a Constant Systems cell disruptor at 25 kpsi. Cell lysates were then clarified by centrifugation at 25,000 g for 20 min at 4 °C. VLPs were isolated from the clarified cell lysates by loading on an iodixanol gradient (15, 20, 25% v/v) and centrifugation at 150,000 g for 3 h at 4 °C using a Beckman Coulter L-100XP floor standing ultracentrifuge. VLP protein pellets were then further purified by size exclusion chromatography (SEC) on a HiPrep Sephacryl 16/60 S-500 HR column (Cytiva) using an ÄKTA FPLC system. Protein species from SDS-PAGE were identified by in-gel tryptic digest liquid LC-MS/MS on a Sciex 5600 Quadrupole time-of-flight mass spectrometer (QTOF-MS). Protein concentrations were calculated using the Beer-Lambert law from absorbance values obtained using a NanoDrop (Thermofisher).

### AEP Expression, Purification and Activation

The enzyme was expressed in *E. coli* SHuffle cells using a pHUE construct encoding 6His–ubiquitin–[C247A]*Oa*AEP1 as described previously^53^. Expression was induced with 0.4 mM IPTG with overnight incubation at 18 °C. Cells were lysed and the protein purified from clarified lysates by Ni–NTA affinity chromatography prior to desalting into activation buffer. Auto-activation was initiated by lowering the pH to 4.0–4.5 with acetic acid, followed by incubation at 37 °C for 1 h, with activation monitored by SDS–PAGE. Protein concentration was determined as described for VLP substrates, and aliquots were stored at −80 °C.

### Production of Protein Nucleophiles

For expression of both GL-sfGFP, GVG-ZHER2, and GVG-9G8, BL21(DE3) *E. coli* cultures were induced at A600 of 0.6 with 1 mM IPTG and incubated overnight at 18 °C with shaking at 220 rpm. Cells, harvested by centrifugation at 4,500 g for 15 min at 4 °C, were resuspended in binding buffer (500 mM NaCl, 50 mM Tris-HCl, 20 mM imidazole, pH 8 containing EDTA-free protease inhibitor) before passage through a Constant Systems cell disruptor at 25 kpsi. Cell lysates were clarified by centrifugation at 40,000 g for 30 min at 4 °C, passed through a 0.45 µm filter and then loaded onto a HisTrap FF column (Cytiva) for purification using an ÄKTA FPLC system. Fractions from the HisTrap purification were analyzed by SDS-PAGE and concentrated using an Amicon filter. GL-sfGFP was cleaved with Factor Xa (New England Biolabs) and GVG-ZHER2 and GVG-9G8 were cleaved with hyperTEV60 as per manufacturer guidelines and the published protocol^54^. Protein concentrations were determined as described for other constructs.

### Synthesis and Purification of GVG-PDIP

GVG-PDIP was synthesized from C- to N-termini on 2-chlorotitryl resin using automated Fmoc solid phase chemistry (Symphony, Peptide Technologies Inc.). The synthesized peptide was removed from resin with simultaneous removal of protecting groups by adding trifluoroacetic acid (TFA), water and triisopropylsilane (TIPS) (95:2.5:2.5 v/v/v) for 2.5 h before inducing phase separation of peptide from TIPS/TFA by addition of ice-cold diethyl ether 45% (v/v) acetonitrile (ACN), 0.05% (v/v) TFA. Residual ether/TFA was evaporated on a Rotary Evaporator (40 °C, 200 mBar) for 20 min before overnight lyophilization. The cleaved peptide mixture was purified by reverse-phase high-performance liquid chromatography (RP-HPLC) using a Shimadzu preparative system and Phenomenex Jupiter C18 column with a 1% per min gradient of solvent B (90% v/v ACN, 0.05% v/v TFA) against solvent A (0.05% v/v TFA). Purity and correct mass were simultaneously determined using a Shimadzu LCMS-2020 instrument with a Phenomenex 5 µm C18 / 300 Å / 150 x 2 mm LC column. (Figure S1). Chemical synthesis, purification, and purity evaluation of synthetic GVG-PDIP were performed by Dr Crystal Huang and Mr Lachlan Hall (University of Queensland).

### Negative Stain Transmission Electron Microscopy

Purified VLPs were diluted to 0.25 mg/mL in PBS and applied (10 µL) to formvar/carbon coated grids (ProSciTech) for 2 min. Grids were washed three times on droplets of water for 30 s before staining on 10 µL of 2 % (v/v) uranyl acetate for 2 min. TEM was performed on a JEOL 1400 electron microscope at 80 kV operating voltage.

### AEP-Mediated Transpeptidation Reactions

Reactions for characterization of transpeptidation for all tested VLP-conjugates were conducted in 30 µL PBS (pH 7.4) containing 15 µM VLP (calculated concentration of CP-NGLH), 60 µM conjugation partner, 1 µM AEP, and 150 µM NiSO_4_. Time point reaction quenching was achieved by taking an aliquot and diluting 1:1 in 1% (v/v) formic acid before assessment by SDS-PAGE. For scaled up reactions, conditions were kept the same except at increased volumes (up to 1 mL) and reactions were terminated by repeated dilution and washing through a 100 kDa Amicon filter to separate AEP and remaining unreacted conjugation partners from functionalized VLPs.

### Production of Recombinant Receptors

Gene coding sequences for Human HER2 extracellular domain (UniProt ID 04626, T23-T652) with a C-terminal StrepII tag and EGFR extracellular domain (UniProt ID P00533, L25-S645) with a C-terminal 6xHis tag, both containing N-terminal Immunoglobulin Kappa signal peptide and 5’ kozak sequence, were synthesized and cloned into pTwist CMV BetaGlobin WPRE Neo by Twist Bioscience. Plasmid DNA was transfected into CHO cells using the ExpiCHO expression system (Gibco) and cultured following suppliers recommended protocol for maximum titer expression. Cell cultures were then harvested by centrifugation and supernatants filtered at 0.45 µm (Millipore) before affinity chromatography. HER2 was purified using a 5 ml StrepTrap XT column (Cytiva) while EGFR was purified using a 5 ml HisTrap excel column (Cytiva) both following supplier’s column recommendations. Purity was evaluated by SDS-PAGE on a 4-12% gradient Bolt Bis-Tris gel (Invitrogen) and concentration determined by UV-Vis using the N120 NanoPhotometer (Implen).

### Receptor Binding Assay

Black 96-well microtiter plate was coated overnight in recombinant receptor (HER2 or EGFR) at 5 µg/mL or 1% (w/v) bovine serum albumin (BSA) in 20 mM Tris-HCl (pH 7.4), 150 mM NaCl, 2 mM CaCl_2_, 1 mM MnCl_2_, and 1 mM MgCl_2_. Wells were washed 3 times with PBS (pH 7.4) before blocking with 1% BSA for 90 min at 37 °C. Wells were washed again 3 times with PBS before incubation with VLPs at 5 µg/mL for 30 min at 37 °C. Following another 3 washes in PBS, fluorescence intensity was recorded on a TECAN SparkControl plate reader with excitation at 540 nm and emission recorded at 592 nm. Background fluorescence of receptors was subtracted, and data normalized to non-specific VLP binding to BSA alone, with 5 technical replicates recorded for each construct. Statistical significance was determined by a Welch’s t-test using GraphPad Prism v10.

## AUTHOR INFORMATION

### Author Contributions

**Maxim D. Harding** conceptualization, data curation, formal analysis, investigation, methodology, project administration, validation, visualization, writing – original draft; **Mark A. Jackson** funding acquisition, methodology, resources, supervision, writing – review and editing; **Kuok Yap** investigation, methodology, writing – review and editing; **Pie Huda** resources, writing – review and editing; **David J. Craik** funding acquisition, resources, supervision, writing – review and editing; **Frank Sainsbury** conceptualization, methodology, resources, validation, writing – review and editing; **Nicole Lawrence** funding acquisition, methodology, project administration, resources, supervision, validation, writing – review and editing.

### Notes

Authors declare no conflicts of interest.

## Acknowledgements

The Authors would like to acknowledge Dr Crystal Huang and Mr Lachlan Hall for help with peptide synthesis and purification as well as Dr Yan Zhou for providing the GFP construct. The authors gratefully acknowledge the support and infrastructure provided by the National Biologics Facility.

## ASSOCIATED CONTENT

### Supporting Information

The following files are available free of charge.

Amino acid sequences of proteins used in this study; tryptic digest LC-MS/MS coverage of recombinant proteins; characterization of synthetic GVG-PDIP; western blot analysis of CP-PDIP conjugate; VLP aggregation upon 16 h conjugation with GVG-PDIP; recombinant GL-sfGFP; SDS-PAGE of GL-sfGFP conjugation replicates; P22 VLP hexamer structure; recombinant GVG-ZHER2; recombinant GVG-9G8; SDS-PAGE of GVG-ZHER2 and GVG-9G8 conjugation replicates (PDF).

## Entry for Table of Contents

We employ an asparaginyl endopeptidase (AEP) ligase for late-stage functionalization of virus-like particles (VLPs) from bacteriophage P22. AEP-mediated conjugation of peptides and proteins onto assembled P22 VLPs is extremely efficient, requiring low enzyme concentration and minimal excess of nucleophile, in a short time. VLPs functionalized with receptor-biding domains bind to cognate receptors *in vitro*.

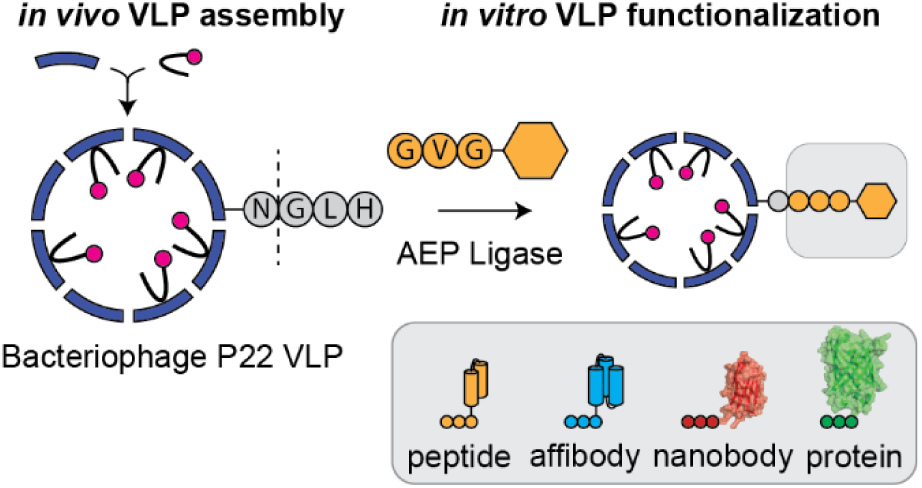

