## Supporting Information for "Selective and Efficient Functionalization of P22 Virus-Like Particles Using an Asparaginyl Ligase"

**Contents**

Table S1. Amino acid sequences of proteins used in this study.

Table S2. Tryptic digest LC-MS/MS of recombinant proteins.

Figure S1. Characterization of synthetic GVG-PDIP.

Figure S2. Western blot analysis of CP-PDIP conjugate.

Figure S3. VLP aggregation upon 16 h conjugation with GVG-PDIP.

Figure S4. Recombinant GL-sfGFP.

Figure S5. SDS-PAGE of GL-sfGFP conjugation replicates.

Figure S6. P22 VLP hexamer structure.

Figure S7. Recombinant GVG-ZHER2.

Figure S8. Recombinant GVG-9G8.

Figure S9. SDS-PAGE of GVG-ZHER2 and GVG-9G8 conjugation replicates.

**Table S1. Amino acid sequences of proteins used in this study.**

| Protein | Sequence [a] |
|---|---|
| CP-<br>NGLH<br>47.48<br>kDa | MALNEGQIVTLAVDEIIETISAITPMAQKAKKYTPPAASMQRSSNTIWMP<br>VEQESPTQEGWDLTDKATGLLELNVAVNMGEPDNDFFQLRADDLRDETAY<br>RRRIQSAARKLANNVELKVANMAAEMGSLVITSPDAIGTNTADAWNFVAD<br>AEEIMFSRELNRDMGTSYFFNPQDYKKAGYDLTKRDIFGRIPEEAYRDGT<br>IQRQVAGFDDVLRSPKLPVLTKSTATGITVSGAQSFKPVAWQLDNDGNKV<br>NVDNRFATVTLSATTGMKRGDKISFAGVKFLGQMAKNVLAQDATFSVVRV<br>VDGTHVEITPKPVALDDVSLSPEQRAYANVNTSLADAMAVNILNVKDART<br>NVFWADDAIRIVSQPIPANHELFAGMKTTSFSIPDVGLNGIFATQGDIST<br>LSGLCRIALWYGVNATRPEAIGVGLPGQTA**GGSGGNGLH** |
| SP-<br>mRUBY3<br>35.31<br>kDa | MTRLSERLTLKPRGKQISSAPPADQPITGDVSAANKDAIRKQMDAAASKG<br>DVETYRKLKAKLKGIRGGGGSGGGGSMVSKGEELIKENMRMKVVMEGSVN<br>GHQFKCTGEGEGRPYEGVQTMRIKVIEGGPLPFAFDILATSFMYGSRTFI<br>KYPADIPDFFKQSFPEGFTWERVTRYEDGGVVTVTQDTSLEDGELVYNVK<br>VRGVNFPSNGPVMQKKTKGWEPNTEMMYPADGGLRGYTDIALKVDGGGHL<br>HCNFVTTYRSKKTVGNIKMPGVHAVDHRLERIEESDNETYVVQREVAVAK<br>YSNLGGGMDELYKGSHHHHHH |
| GL-<br>sfGFP<br>28.94<br>kDa | MGLSRKGEELFTGVVPILVELDGDVNGHKFSVRGEGEGDATNGKLTLKFI<br>CTTGKLPVPWPTLVTTLTYGVQCFARYPDHMKQHDFFKSAMPEGYVQERT<br>ISFKDDGTYKTRAEVKFEGDTLVNRIELKGIDFKEDGNILGHKLEYNFNS<br>HNVYITADKQKNGIKANFKIRHNVEDGSVQLADHYQQNTPIGDGPVLLPD<br>NHYLSTQSVLSKDPNEKRDHMVLLEFVTAAGITHGMAELYKGSGALGSGI<br>EGRGHHHHHH |
| GVG-<br>ZHER2<br>9.37 kDa | MHHHHHHHHGGENLYFQGVGGSGGGVDNKFNKEMRNAYWEIALLPNLNNQ<br>QKRAFIRSLYDDPSQSANLLAEAKKLNDAQAPK |
| GVG-<br>9G8<br>16.78<br>kDa | MSHHHHHHHHGGENLYFQGVGGSGGGEVQLVESGGGLVQAGGSLRLSCAA<br>SGRTFSSYAMGWFRQAPGKEREFVVAINWSSGSTYYADSVKGRFTISRDN<br>AKNTMYLQMNSLKPEDTAVYYCAAGYQINSGNYNFKDYEYDYWGQGTQVT<br>VSS |

[a] The additional 9 amino acids appended the C-terminus of P22 CP for AEP recognition are highlighted in **bold,** with the optimized AEP-recognition sequence **underlined**.

For proteins GL-sfGFP, GVG-ZHER2, and GVG-9G8, the final protein sequences after cleavage with Factor Xa and TEV are underlined, respectively. The methionine of GL-sfGFP is cleaved *in vivo* in *E. coli* by methionine aminopeptidase.

**Table S2. Tryptic digest LC-MS/MS of recombinant proteins.**

| Protein | Sequence [a] |
|---|---|
| CP-NGLH | MALNEGQIVTLAVDEIIETISAITPMAQKAKKYTPPAASMQRSSNTIWMP<br>VEQESPTQEGWDLTDKATGLLELNVAVNMGEPDNDFFQLRADDLRDETAY<br>RRRIQSAARKLANNVELKVANMAAEMGSLVITSPDAIGTNTADAWNFVAD<br>AEEIMFSRELNRDMGTSYFFNPQDYKKAGYDLTKRDIFGRIPEEAYRDGT<br>IQRQVAGFDDVLRSPKLPVLTKSTATGITVSGAQSFKPVAWQLDNDGNKV<br>NVDNRFATVTLSATTGMKRGDKISFAGVKFLGQMAKNVLAQDATFSVVRV<br>VDGTHVEITPKPVALDDVSLSPEQRAYANVNTSLADAMAVNILNVKDART<br>NVFWADDAIRIVSQPIPANHELFAGMKTTSFSIPDVGLNGIFATQGDIST<br>LSGLCRIALWYGVNATRPEAIGVGLPGQTAGGSGGNGLH |
| SP-mRUBY3 | MTRLSERLTLKPRGKQISSAPPADQPITGDVSAANKDAIRKQMDAAASKG<br>DVETYRKLKAKLKGIRGGGGSGGGGSMVSKGEELIKENMRMKVVMEGSVN<br>GHQFKCTGEGEGRPYEGVQTMRIKVIEGGPLPFAFDILATSFMYGSRTFI<br>KYPADIPDFFKQSFPEGFTWERVTRYEDGGVVTVTQDTSLEDGELVYNVK<br>VRGVNFPSNGPVMQKKTKGWEPNTEMMYPADGGLRGYTDIALKVDGGGHL<br>HCNFVTTYRSKKTVGNIKMPGVHAVDHRLERIEESDNETYVVQREVAVAK<br>YSNLGGGMDELYKGSHHHHHH |
| CP-NGL-sfGFP | MALNEGQIVTLAVDEIIETISAITPMAQKAKKYTPPAASMQRSSNTIWMP<br>VEQESPTQEGWDLTDKATGLLELNVAVNMGEPDNDFFQLRADDLRDETAY<br>RRRIQSAARKLANNVELKVANMAAEMGSLVITSPDAIGTNTADAWNFVAD<br>AEEIMFSRELNRDMGTSYFFNPQDYKKAGYDLTKRDIFGRIPEEAYRDGT<br>IQRQVAGFDDVLRSPKLPVLTKSTATGITVSGAQSFKPVAWQLDNDGNKV<br>NVDNRFATVTLSATTGMKRGDKISFAGVKFLGQMAKNVLAQDATFSVVRV<br>VDGTHVEITPKPVALDDVSLSPEQRAYANVNTSLADAMAVNILNVKDART<br>NVFWADDAIRIVSQPIPANHELFAGMKTTSFSIPDVGLNGIFATQGDIST<br>LSGLCRIALWYGVNATRPEAIGVGLPGQTAGGSGGNGGLSRKGEELFTGV<br>VPILVELDGDVNGHKFSVRGEGEGDATNGKLTLKFICTTGKLPVPWPTLV<br>TTLTYGVQCFARYPDHMKQHDFFKSAMPEGYVQERTISFKDDGTYKTRAE<br>VKFEGDTLVNRIELKGIDFKEDGNILGHKLEYNFNSHNVYITADKQKNGI<br>KANFKIRHNVEDGSVQLADHYQQNTPIGDGPVLLPDNHYLSTQSVLSKDP<br>NEKRDHMVLLEFVTAAGITHGMAELYKGSGALGSGIEGR |
| CP-NGVG-ZHER2 | MALNEGQIVTLAVDEIIETISAITPMAQKAKKYTPPAASMQRSSNTIWMP<br>VEQESPTQEGWDLTDKATGLLELNVAVNMGEPDNDFFQLRADDLRDETAY<br>RRRIQSAARKLANNVELKVANMAAEMGSLVITSPDAIGTNTADAWNFVAD<br>AEEIMFSRELNRDMGTSYFFNPQDYKKAGYDLTKRDIFGRIPEEAYRDGT<br>IQRQVAGFDDVLRSPKLPVLTKSTATGITVSGAQSFKPVAWQLDNDGNKV<br>NVDNRFATVTLSATTGMKRGDKISFAGVKFLGQMAKNVLAQDATFSVVRV<br>VDGTHVEITPKPVALDDVSLSPEQRAYANVNTSLADAMAVNILNVKDART<br>NVFWADDAIRIVSQPIPANHELFAGMKTTSFSIPDVGLNGIFATQGDIST<br>LSGLCRIALWYGVNATRPEAIGVGLPGQTAGGSGGNGVGGSGGGVDNKFN<br>KEMRNAYWEIALLPNLNNQQKRAFIRSLYDDPSQSANLLAEAKKLNDAQA<br>PK |
| CP-NGVG-9G8 | MALNEGQIVTLAVDEIIETISAITPMAQKAKKYTPPAASMQRSSNTIWMP<br>VEQESPTQEGWDLTDKATGLLELNVAVNMGEPDNDFFQLRADDLRDETAY<br>RRRIQSAARKLANNVELKVANMAAEMGSLVITSPDAIGTNTADAWNFVAD<br>AEEIMFSRELNRDMGTSYFFNPQDYKKAGYDLTKRDIFGRIPEEAYRDGT<br>IQRQVAGFDDVLRSPKLPVLTKSTATGITVSGAQSFKPVAWQLDNDGNKV<br>NVDNRFATVTLSATTGMKRGDKISFAGVKFLGQMAKNVLAQDATFSVVRV |

| | |
|---|---|
| | VDGTHVEITPKPVALDDVSLSPEQRAYANVNTSLADAMAVNILNVKDART<br>NVFWADDAIRIVSQPIPANHELFAGMKTTSFSIPDVGLNGIFATQGDIST<br>LSGLCRIALWYGVNATRPEAIGVGLPGQTAGGSGGNGVGGSGGGEVQLVE<br>SGGGLVQAGGSLRLSCAASGRTFSSYAMGWFRQAPGKEREFVVAINWSSG<br>STYYADSVKGRFTISRDNAKNTMYLQMNSLKPEDTAVYYCAAGYQINSGN<br>YNFKDYEYDYWGQGTQVTVSS |

[a] Underlined sections represent peptide sequences identified by in-gel tryptic digest LC-MS/MS analysis with high confidence.

**A)**

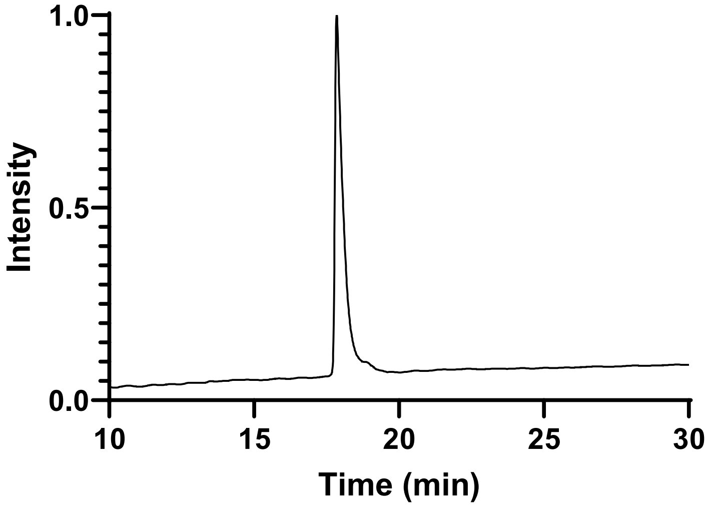


**B)** Calculated mass (oxidized): 3989.84 Da Observed mass: 3988.8 Da

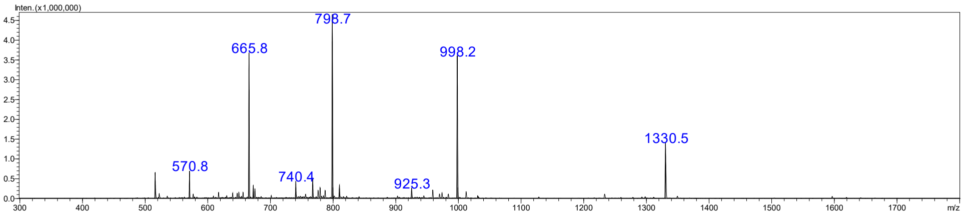


**Figure S1. Characterization of synthetic GVG-PDIP. A)** LC trace of purified synthetic GVG-PDIP produced for this study. **B)** Electrospray ionization mass spectrometry signal for purified GVG-PDIP. The calculated molecular weight for oxidized GVG-PDIP is shown along with the observed mass extrapolated from the $[(m+z)/z]^{z+}$ peaks.

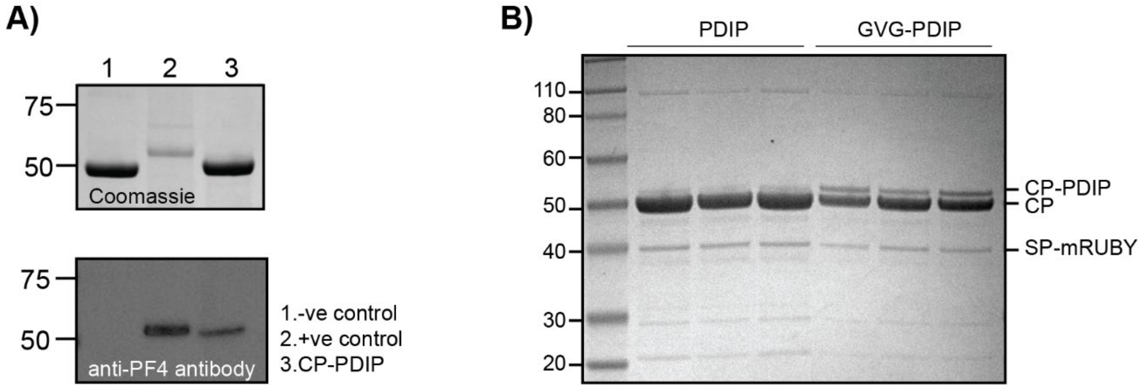


**Figure S2. Western blot analysis of CP-PDIP conjugate. A)** SDS-PAGE and western blot analysis of PDIP conjugation to P22 VLP. Top panel shows Coomassie stained SDS-PAGE gel, bottom panel shows western blot stained with a PDIP-specific antibody (platelet factor 4 polyclonal antibody, Abcam). Lane 1 is a P22 VLP control. Lane 2 is a PDIP-CP fusion, made in *Nicotiana benthamiana*[1]. Lane 3 is P22 CP-PDIP. **B)** Coomassie stained SDS-PAGE gel showing (n=3) 16 h conjugation reactions with either PDIP or GVG-PDIP. Protein species are depicted on right.

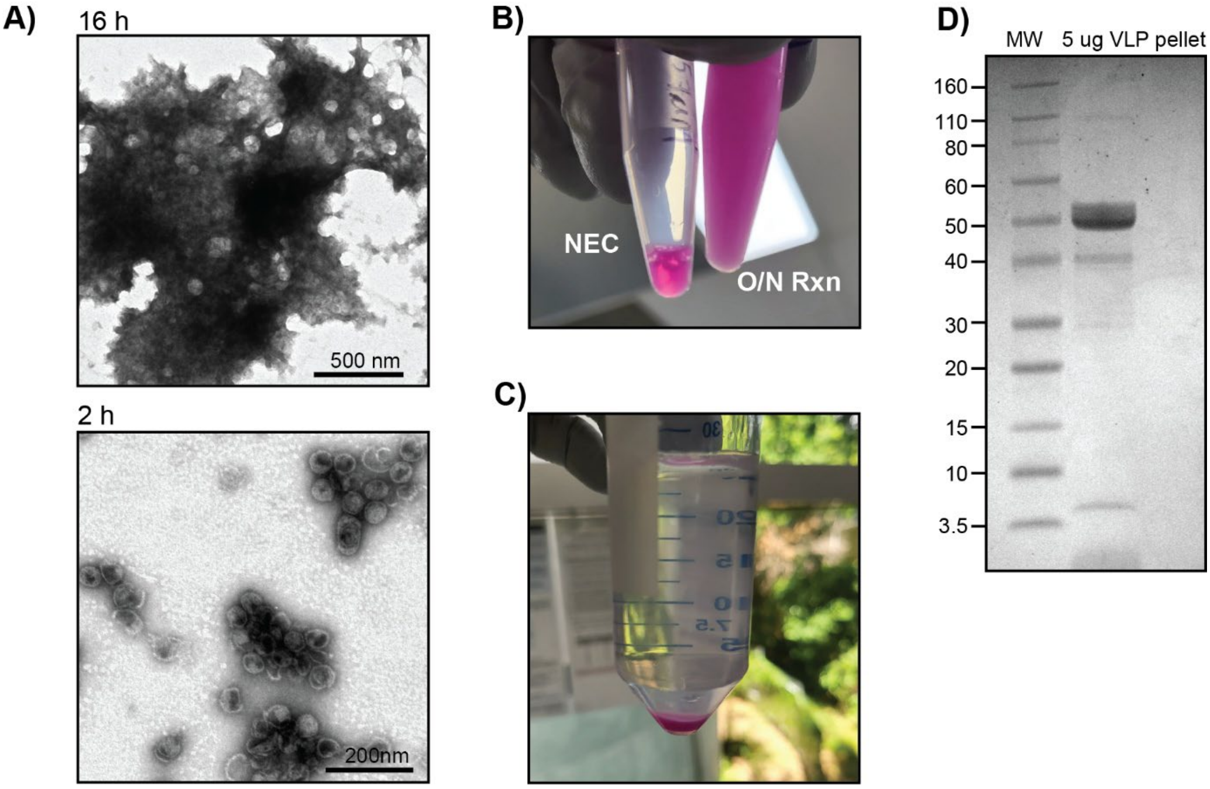


**Figure S3. VLP aggregation upon 16 h conjugation with GVG-PDIP. A)** Negative stain TEM following 16 h or 2 h reaction incubation (note difference in scale bars). **B)** Comparison in reaction turbidity between no enzyme control (NEC) and overnight reaction (O/N Rxn, 16 h). **C)** Pink pellet of aggregated VLPs following dilution of 16 h reaction in 25 mL PBS and centrifugation at 3,900 *g* for 5 min. **D)** Coomassie stained SDS-PAGE gel of resuspended VLP pellet shown in (C).

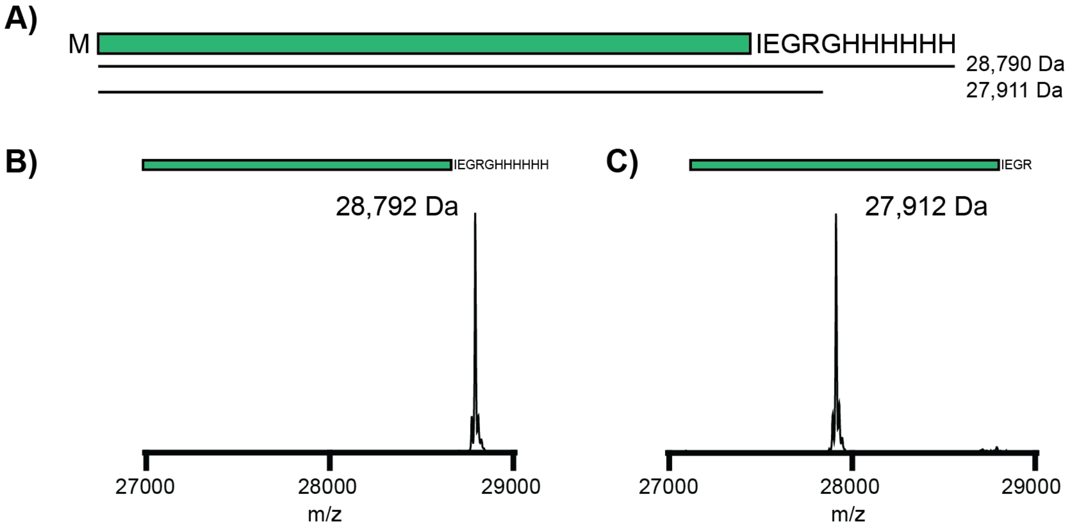


**Figure S4. Recombinant GL-sfGFP. A)** Cartoon of the GL-sfGFP sequence expressed in *E. coli*. The N-terminal methionine is processed endogenously by methionine aminopeptidase, leaving an N-terminal GL (predicted molecular weight 28,790 Da). The C-terminal 6His tag, used for HisTrap purification, is cleaved by Factor Xa following purification (predicted molecular weight 27,911 Da). **B)** Mass of GL-sfGFP following HisTrap purification. **C)** Mass of GL-sfGFP purified following Factor Xa cleavage.

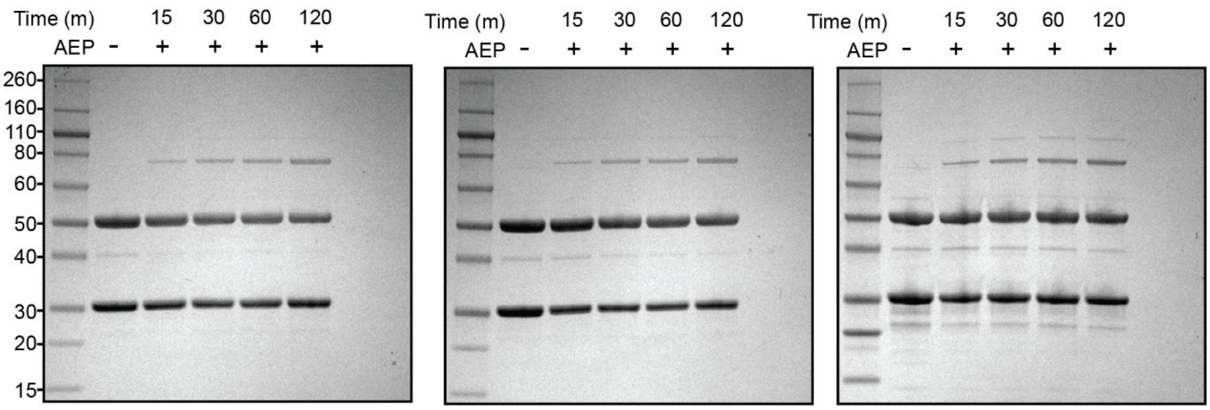


**Figure S5. SDS-PAGE of GL-sfGFP conjugation replicates**. Coomassie stained SDS-PAGE gels from n=3 technical replicates of GL-sfGFP conjugation to P22 VLPs. Time points are shown in minutes above sample lanes. The first lane contains protein ladder (marked in kDa on left side of first gel). The second lane is a no-enzyme control.

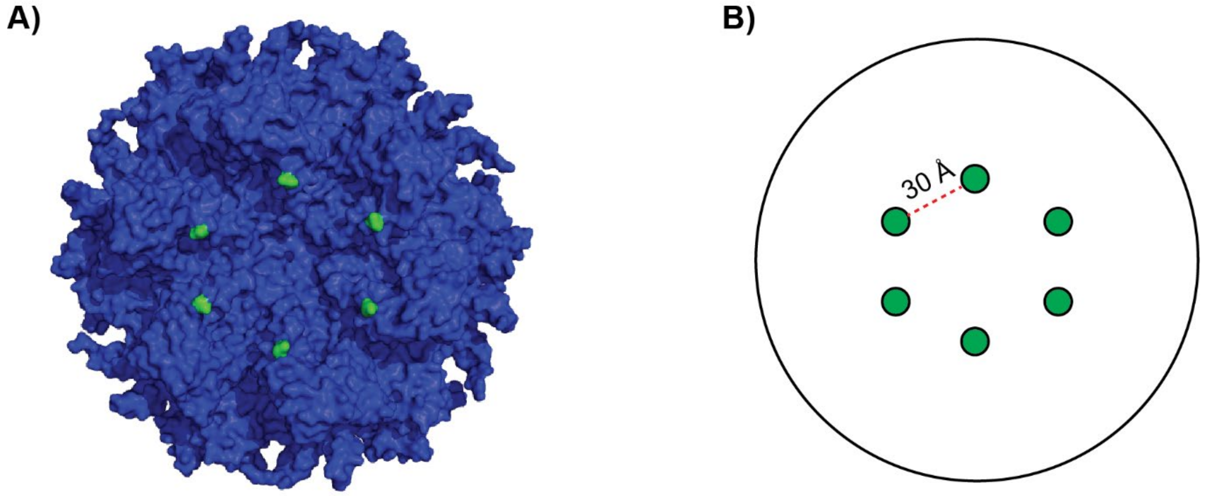


**Figure S6. P22 VLP hexamer structure. A)** Surface illustration of P22 CP hexamer from (PDB: 5UU5). The C-terminal amino acid is highlighted in green, illustrating proximity around the hexameric interface. **B)** An outline of the CP hexamer, illustrating the 30 Å distance between C-termini in adjacent CP monomers at the hexameric interface. Surface image and measurements generated using PyMOL v2.5.2.

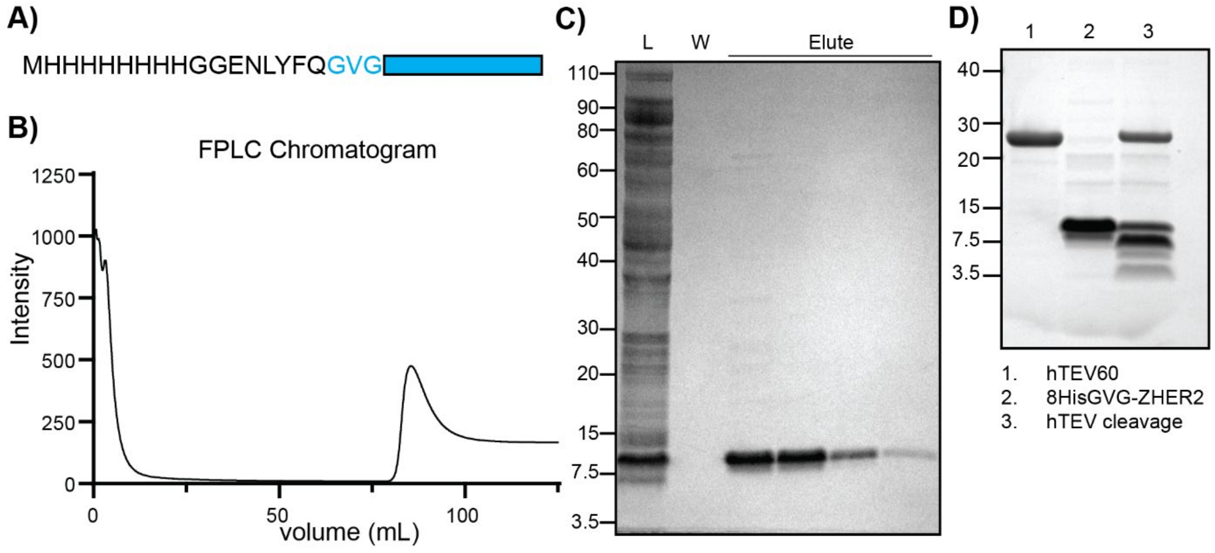


**Figure S7. Recombinant GVG-ZHER2. A)** Cartoon of 8HisGVG-ZHER2 construct, containing an 8His tag and TEV processing site (ENLYFQ). **B)** FPLC chromatogram from HisTrap purification of 8HisGVG-ZHER2 from *E. coli* cell lysate. The peak at ~80 mL is the elution peak containing pure protein. **C)** Coomassie stained SDS-PAGE of the HisTrap purification. The first lane (L) contains total crude protein, second lane (W) is the wash lane, following lanes (elute) are the elution fractions containing purified 8HisGVG-ZHER2. **D)** Coomassie stained SDS-PAGE showing (1) hTEV60, (2) purified 8HisGVG-ZHER2, and (3) product of incubation of both hTEV60 and 8HisGVG-ZHER2 overnight, with the TEV cleavage product of GVG-ZHER2 shown at 7.5 kDa.

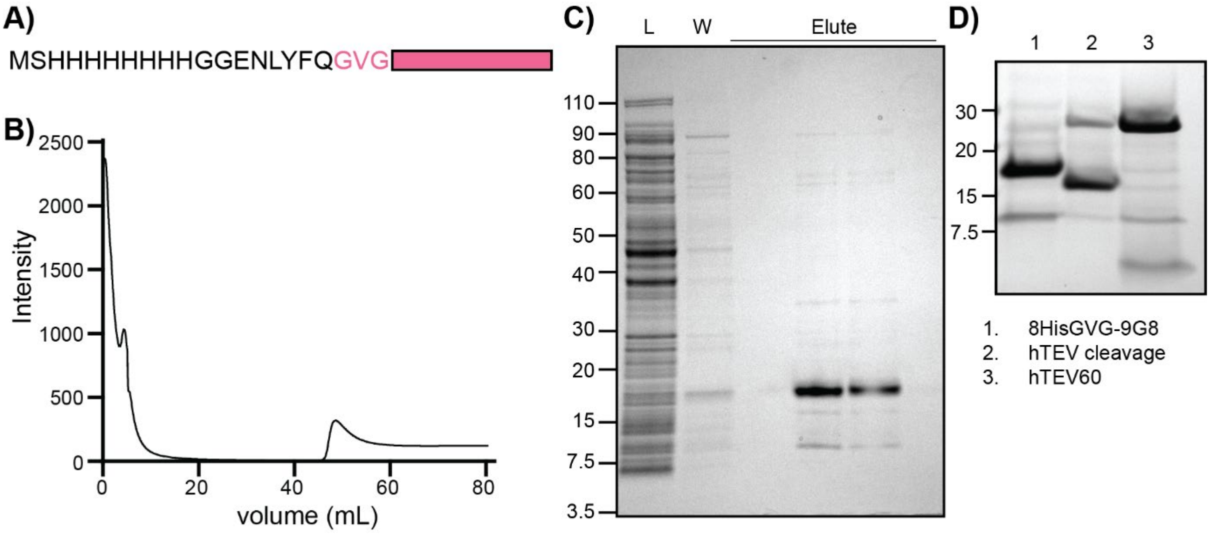


**Figure S8. Recombinant GVG-9G8. A)** Cartoon of 8HisGVG-9G8 construct, containing an 8His tag and TEV processing site (ENLYFQ). **B)** FPLC chromatogram from HisTrap purification of 8HisGVG-9G8 from *E. coli* cell lysate. The peak at ~50 mL is the elution peak containing pure protein. **C)** Coomassie stained SDS-PAGE of the HisTrap purification. The first lane (L) contains total crude protein, second lane (W) is the wash lane, following lanes (elute) are the elution fractions containing purified 8HisGVG-9G8. **D)** Coomassie stained SDS-PAGE showing (1) purified 8HisGVG-9G8, (2) product of incubation of both hTEV60 and 8HisGVG-9G8 overnight, with the TEV cleavage product of GVG-9G8 shown at approximately 16 kDa and (3) hTEV60 alone.

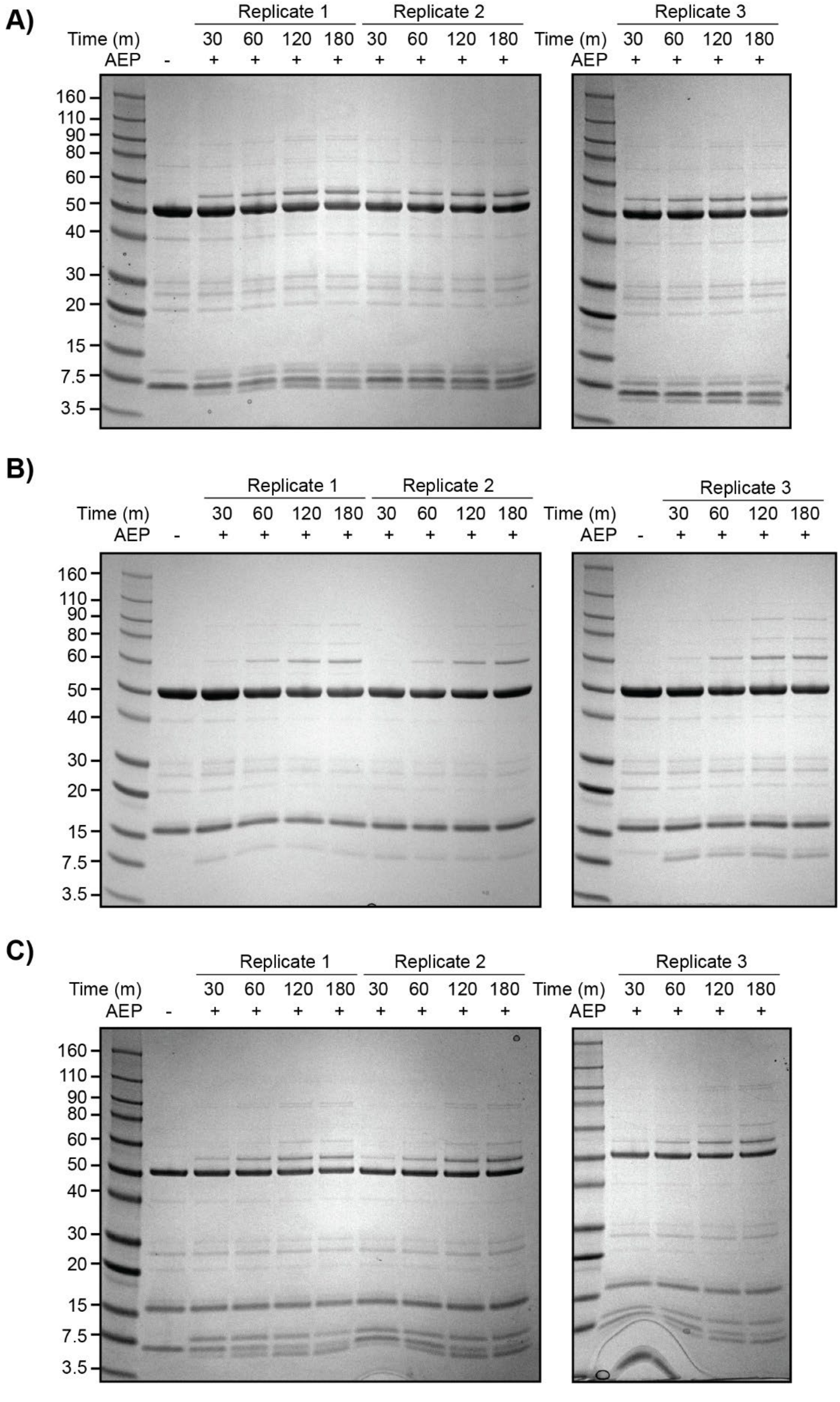


**Figure S9. SDS-PAGE of GVG-ZHER2 and GVG-9G8 conjugation replicates. A)** Conjugation with GVG-ZHER2. **B)** Conjugation with GVG-9G8. **C)** One-pot dual conjugation reaction with both GVG-ZHER2 and GVG-9G8.
